# Open-Source Tools to Analyze Temporal and Spatial Properties of Local Field Potentials

**DOI:** 10.1101/2024.03.14.584529

**Authors:** Geoffrey M. Barrett, Srujan Vajram, Oliver Shetler, Andrew Aoun, S. Abid Hussaini

**Affiliations:** Taub Institute for Research on Alzheimer’s disease and the Aging Brain, Columbia University Irving Medical Center, New York, NY, 10032, USA; Department of Pathology and Cell Biology, Columbia University Irving Medical Center, New York, NY, 10032, USA

## Abstract

Analysis of local field potentials (LFPs) is important for understanding how ensemble neurons function as a network in a specific region of the brain. Despite the availability of tools for analyzing LFP data, there are some missing features such as analysis of high frequency oscillations (HFOs) and spatial properties. In addition, accessibility of most tools is restricted due to closed source code and/or high costs. To overcome these issues, we have developed two freely available tools that make temporal and spatial analysis of LFP data easily accessible. The first tool, hfoGUI (High Frequency Oscillation Graphical User Interface), allows temporal analysis of LFP data and scoring of HFOs such as ripples and fast ripples which are important in understanding memory function and neurological disorders. To complement the temporal analysis tool, a second tool, SSM (Spatial Spectral Mapper), focuses on the spatial analysis of LFP data. The SSM tool maps the spectral power of LFPs as a function of subject’s position in a given environment allowing investigation of spatial properties of LFP signal. Both hfoGUI and SSM are open-source tools that have unique features not offered by any currently available tools, and allow visualization and spatio-temporal analysis of LFP data.

## Introduction

Investigating the brain’s network activity is important for understanding how populations of neurons communicate with each other and how this relates to cognitive function. Local field potentials (LFPs) are oscillations resulting from the firing of many neurons in a specific region of the brain. Unlike the electroencephalogram (EEG), which are collected from a standard set of positions from the surface of a skull, LFPs are collected using depth electrodes positioned in one or more regions of the brain, such as the hippocampus or the entorhinal cortex. EEG is practical for human use because it is non-invasive and is therefore used as the de facto standard for assessing brain health. For example it is routinely used to isolate epileptic seizures. EEG is also used to detect the early onset of neurodegenerative disorders such as Alzheimer’s disease [1–4]. Accordingly, plenty of research focuses on decomposing EEG spectra to examine their role in spatial memory. However, LFPs have superior signal quality because of the electrodes’ proximity to neurons of interest. They are useful for measuring activity of a specific microcircuit users want to understand. Therefore, LFPs are common in rodent research where scientists have a specific hypothesis related to the region of interest. Even though they need different electrodes and have different underlying signal sources, the principles of signal processing applied to both are the same. The signals are often termed as oscillations or rhythms and the tools used for analyzing them are generally interchangeable.

Most tools focus on understanding the temporal properties of brain signals since oscillations of specific frequencies such as delta, theta, gamma, ripples and fast ripples etc. are known to provide an understanding of specific brain function. Delta frequencies (0-3 Hz) are slow brain waves and often used for scoring sleep stages. Theta frequencies (4-12 Hz) are known to be important during animal’s running, grooming as well as memory processing. Gamma frequencies (Low: 35-55 Hz and High: 65-120 Hz) are known to be important for attention and memory. Ripples (80-250 Hz) fall under high frequency oscillations (or HFOs) and are known to be important for memory consolidation. Fast ripples (250-500 Hz) too are part of HFOs and can be linked to either memory consolidation or epilepsy depending on the existence of inter Ictal spikes or epileptiform seizure events in which case they are termed pathological ripples [5–7]. There are several tools available for analyzing LFP data and a recent review highlights their features [8]. However, a majority of these tools are proprietary software, expensive and are not open-source such as MATLAB. Among Python tools, only one of them (Elephant [9]) has LFP analysis but some data types, including our Axona data format, are not compatible and detection of HFOs is not possible.

The analysis of spatial properties of LFPs is key to understand how the brain’s oscillations are influenced by different behaviors. It is important to note that “spatial” in our case refers to a subject’s physical location in an environment. Several researchers do in fact study spatial properties from different EEG electrode locations on the skull which is also important in understanding how various brain regions (mainly cortical) are influenced or co-activated during behavior or affected in disease [10–12]. Since we are interested in how the brain encodes subjects’ physical location in an environment and responds to other navigation-based behaviors, spatial properties of LFPs is of high significance [13, 14]. To our knowledge, unlike tools for analyzing temporal properties of LFP, there are no tools for analyzing spatial properties.

To address the above-mentioned limitations and to overcome the need for analysis of HFOs and spatial component of LFP signal, we have developed two complimentary tools, (a) hfoGUI (High Frequency Oscillations Graphic User Interface), which analyzes temporal properties of LFPs including HFOs and (b) SSM (Spatial Spectral Mapper), which analyzes spatial properties of LFPs. This paper demonstrates that the temporal and spatial properties of local field potentials can be analyzed using free, open-source tools such as hfoGUI and SSM. Despite the availability of numerous software and tools, our tools offer specific advantages not found in others.

## Methods and Results

### HfoGUI— Visualization and Temporal Analysis of LFPs

HfoGUI (High Frequency Oscillation Graphic User Interface) is designed for temporal analysis of frequencies derived from local field potential (LFPs) recordings in mice but potentially any continuous series signal (e.g. EEG) data can be analyzed. HfoGUI is an intuitive, user-friendly tool that can be navigated by a novice with very little training. It boasts several core features including the ability to set various filter-methods, types, and orders, cutoff frequencies, etc. to visualize any number of frequencies simultaneously (Figure 1). It is currently compatible with Axona data format but we provide a tool to convert data from Intan recording system to Axona format which is described in the Data Types section below. The user simply has to pick a file to load and set filters to visualize the frequencies at different time points. HfoGUI is interactive and allows the user to explore detailed frequency information by clicking on a specific timepoint. The T-F Plot Window (Figure 4) allows visualization of time frequency plots using Stockwell transformation along with raw and filtered signals and visualize power spectral density (PSD) plots.

**Figure 1.**
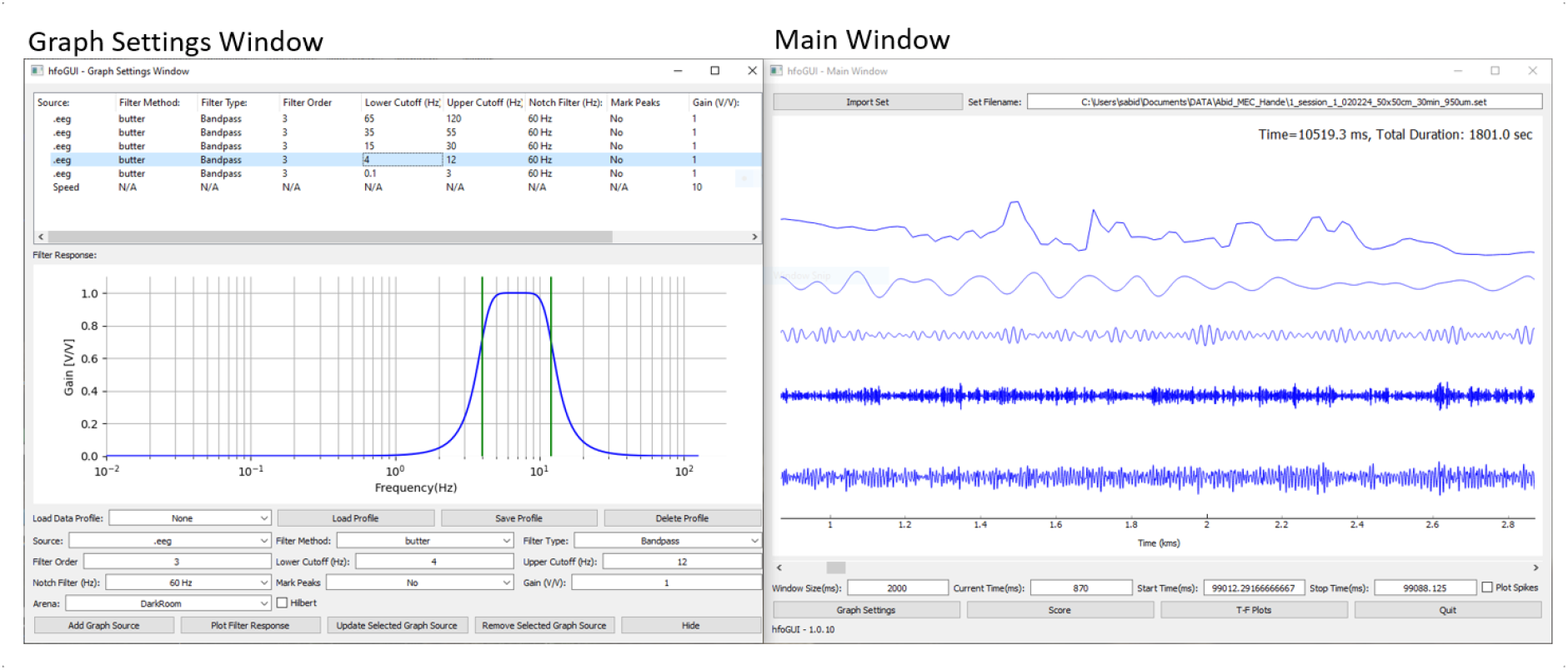
The “Graph Settings” button in the bottom of the Main Window (right) opens the Graph Settings Window (left). To add plots to the Main Window, a source is selected and other parameters are set to visualize frequencies. In the above case theta (4-12Hz), low gamma (35-55Hz), Ripples (80-250Hz) and Fast Ripples (250-500Hz) are the frequencies visualized. In addition, animal’s speed is also added as a source. The bottom panel in the Graph Settings Window shows frequency response of theta band.

HfoGUI allows the user to score and analyze specific events of interest (EOI) within the frequencies in the Score Window. The user can either manually score specific EOIs or use the built-in “automatic detection” method to detect HFOs by setting different parameters. While there are many different HFO detection methods [15], the hfoGUI tool uses the Hilbert transformation method for automatic detection of HFOs. It uses a combination of implementations used in Burnos’s and Ripplelab’s paper with initial detection strategy similar to that of Burnos et. al. and switching to Ripplelab method to detect HFO peaks [16, 17]. Since the Stockwell transform method used by Burnos et. al. to detect HFOs can be computationally intensive and less efficient for real-time analysis, we decided to use Ripplelab’s signal peak threshold and peak number as the basis for HFO detection [16, 17]. This hybrid method allows for faster detection of HFOs without any loss in detection accuracy. It is flexible enough to allow several detection parameters to be set and provides finer control with manual adjustment of events if needed.

### Installation and Usage

HfoGUI was built using Python was specifically tested using Python 3.7. It leverages the PyQt5 cross-platform framework, making it available compatible with all Operating Systems, however it has only been tested on Windows 10. HfoGUI is available from github.com/HussainiLab/hfoGUI. It can be installed using the pip installation method:

python -m pip install hfoGUI . Please refer to the Github page for more details. Once the hfoGUI has been installed, the user can launch the GUI using the following command:

python -m hfoGUI

### Steps for using hfoGUI

#### 1. Adding a session

In the Main Window, a user must first pick a session by selecting a file (Axona’s EEG or EGF file or an Intan file) and then add the plots to visualize. After loading a file, the “Graph Settings” button at the bottom of the Main window will open a Graph Settings Window (Figure 1).

### 2. Adding frequency plots

To add a plot to the Main Window, a user must select a source from the dropdown menu of the Graph Settings Window. For Axona files, options such as .eeg or .egf will be available to select. Generally, the .egf file should be selected since it is sampled at 4.8 kHz, as it has a higher sampling rate than the .eeg files (sampled at 250 Hz). The .egf file will allow the visualization of HFOs since the upper bounds of an HFO is 500 Hz. The Nyquist rate states that the sampling rate must be at least twice the highest frequency present in the signal. Therefore, 250 Hz from the .eeg files will not be sufficient to properly visualize the entire HFO frequency spectrum. If a position file is available in the session, “Speed” can be used as a source for visualization. To use it, the user must also choose an appropriate “Arena” value that corresponds to the session’s data. Each Arena has a varying pixel-per-meter value that will be used to convert the positions from pixels to centimeters. The existing Arena values are specific to Hussaini lab at this time but future versions will allow a PPM value to be entered directly.

### 3. Frequency parameters

In the Graph Settings Window, users will be able to modify this source’s output with a multitude of parameters listed below:

- **Filter Method:** Users have the option to choose between various filters (Butterworth, Chebyshev 1/2, Elliptic, and Bessel). Frequency responses of these filters are plotted in (Figure 2) and the default is Butterworth. In the frequency response, the signals will be normalized to have a gain of 1.
- **Filter Type:** Users can choose between low-pass, high-pass, and band-pass.

**Figure 2.**
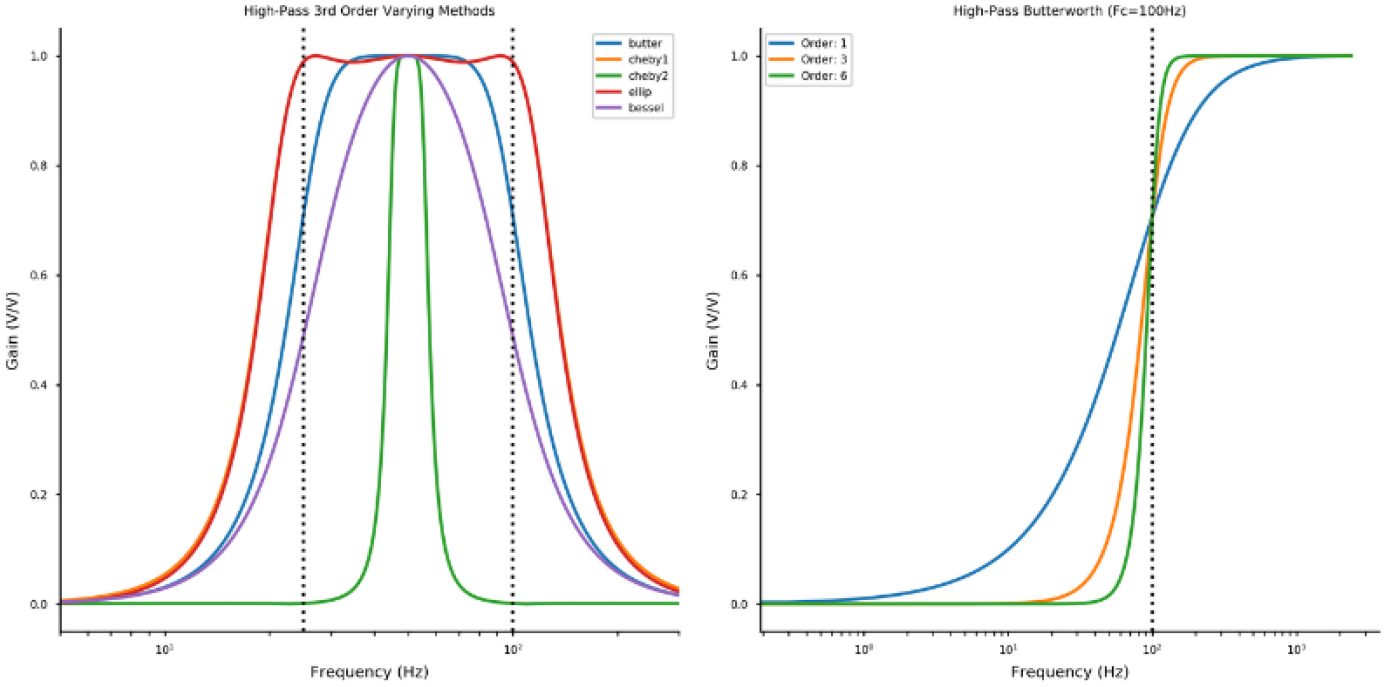
*Left:* Frequency response plots of different filters (Butterworth, Chebyshev 1, Chebyshev 2, Elliptic, and Bessel). Butterworth is the default method used in hfoGUI. *Right:* An example of a high-pass Butterworth filter with different orders plotted to show filter roll-off.

Low-pass filters out frequencies higher than a provided cutoff value, high-pass filters out frequencies below a provided cutoff value, and band-pass filters between two provided cutoff values.

- **Filter Order:** This parameter represents the number of roots/poles of the differential equation describing the filter. This value will affect the filter roll-off (steepness). Some examples of high-pass Butterworth filters with varying orders are plotted just to visualize the impact of order on the frequency response (Figure 2).
- **Lower Cutoff (Hz):** This will be the lower bound cutoff value for a desired filter. For example, 4 would be the lower cutoff value to band-pass filter the signal between 4 Hz and 12 Hz (Theta band).
- **Upper Cutoff (Hz):** This will be the value for the upper bound cutoff for a filter. For a band-pass filter of 4 Hz - 12 Hz, 12 would be the upper cutoff. The maximum value possible for this field is half the sampling rate (Nyquist frequency). Therefore, for a .egf source, the highest upper cutoff value is 2.4 kHz (as it is sampled at 4.8 kHz) and for a .eeg source, the maximum possible value is 125 Hz (as these files are sampled at 250 Hz). Like the lower cutoff, be careful with how close this value is to the Nyquist frequency.
- **Notch Filter (Hz):** The noise from the power supply can be filtered out using notch filtering options: None, 60 Hz, and 50 Hz.
- **Mark Peaks:** This parameter will provide an indication (vertical line) at every local maxima or a peak in the signal (Figure 3). This is useful to align one frequency (e.g., Theta) with another frequency (e.g., Gamma) to determine if the frequencies are aligned or locked with each other. Cross-frequency coupling can be calculated outside of this program.
- **Gain (V/V)**: This parameter will multiply the source’s data by a factor provided in the text field. For example, 2 will double the frequency and 0.5 will halve the frequency.
- **Arena**: This is relevant only if plotting animal speed as a source. The dropdown menu indicates the behavior arena with a specific unit of pixels provided by the camera. At this time only Hussaini lab arenas are supported but an option for entering PPM values will be added soon.
- **Hilbert**: When checked, this parameter will apply the Hilbert Transform to the selected source (after all the filtering and data processing has been performed). The Hilbert transform will produce the Analytic Signal (a signal containing complex-values without the negative frequency components). The Hilbert transform (in red) allows visualization of the envelope of a signal (in blue) and is often used in the automatic detection of HFOs (see Figure 7).

**Figure 3.**
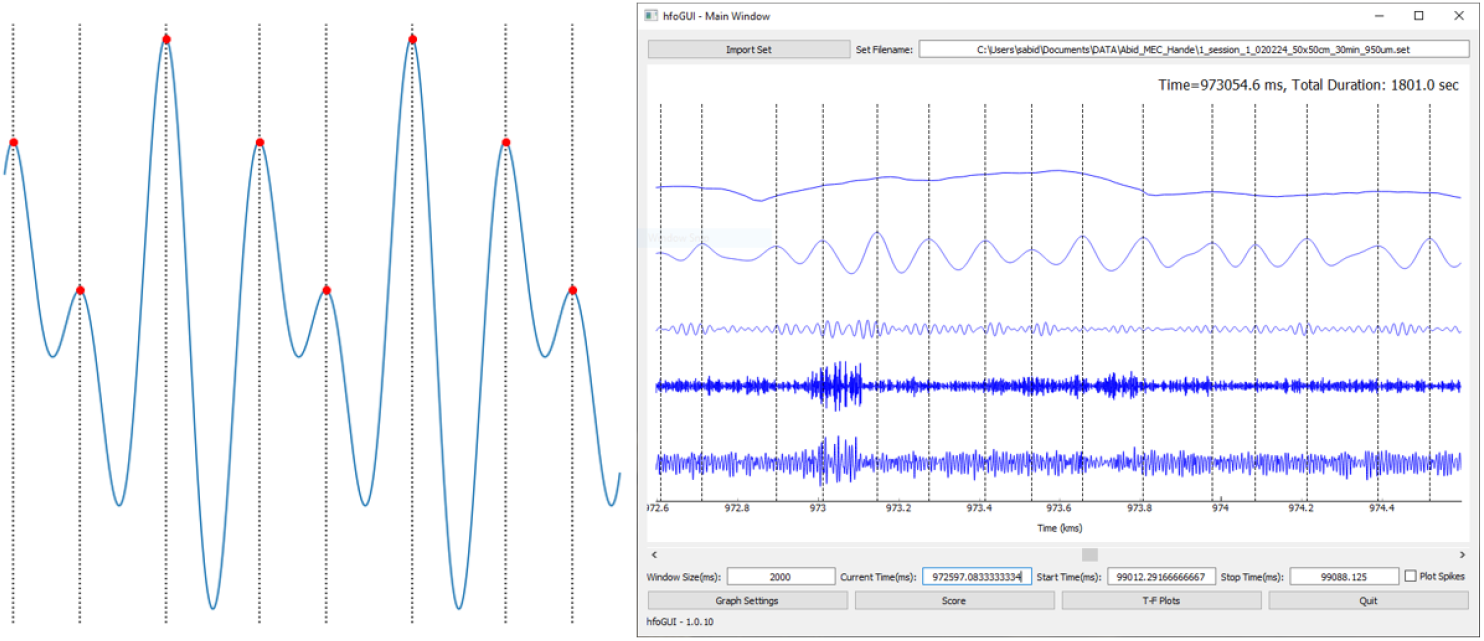
*Mark Frequency Peaks:* When Mark Peaks in the Graph Settings Window for a particular frequency is checked, there is an indication (vertical line) at every local maxima in the signal. Left: The plot shows a close up of the signal and vertical lines (red dots are provided for emphasis).Right: Shows theta frequency (second signal from top) peaks marked with vertical lines. This is useful to quickly count the number of peaks in a signal. In this case one can see 15 peaks in a 2 second window indicating 7.5 Hz signal which confirms the theta frequency band. This is us useful to align one frequency (e.g., Theta) with another frequency (e.g., Gamma) to determine if the frequencies are aligned or locked with each other. Cross-frequency coupling can be calculated outside of this program. Note: This can slow down the visualization if the signal has too many peaks.

**Figure 4.**
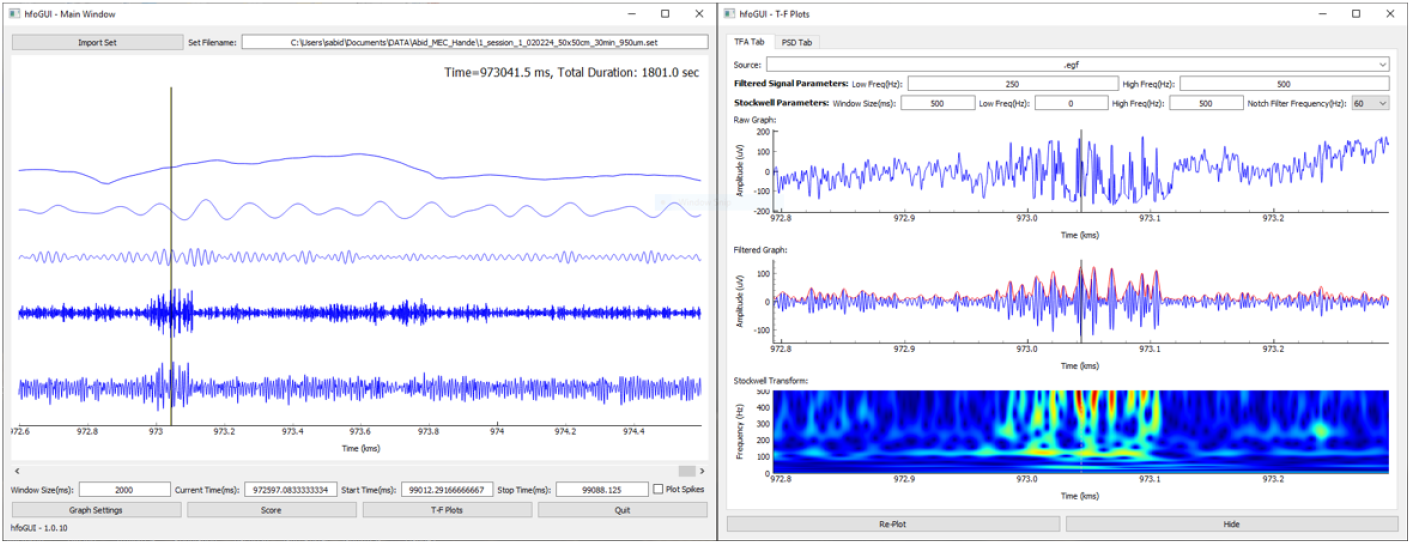
Time-frequency (T-F) Analysis: *Left-* To visualize the T-F of a signal, a user must first click (indicated by a vertical black line) on any specific time point in the Main Window followed by opening the T-F Plots window. *Right-* The default tab that opens in the T-F Plots window is the TFA tab. The top plot shows the raw trace of the selected source, the middle plot is the filtered signal that can be modified by changing the parameters in the Filtered Signal Parameter section at the top of the T-F Plots Window and the bottom plot is the T-F representation using Stockwell transformation centered around the time point that was selected on the Main Window. The heatmap represents red for highest power for a specific frequency (y-axis) at a specific time (x-axis).

**Figure 5.**
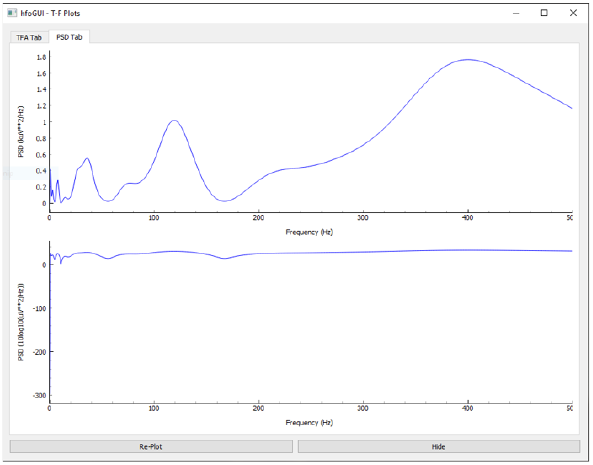
PSD tab: The second tab in the T-F Plots window is the PSD or Power Spectral Density tab. It shows the power of the of the selected signal (vertical line in (Figure 4)) and is represented as V2/Hz. Bottom panel is log PSD.

Multiple sources can be plotted to the Main Window using the “Add Graph Source” button and any plot can be modified by using the “Update Selected Graph Source” button. A set of graph sources can be saved as a profile using the “Save Profile” button and can be quickly loaded the next time a user wants to load them.

### 4. Visualizing Data

In the main window, the graph sources can be visualized at different temporal scales by adjusting the Window Size. This value represents the amount of time (in milliseconds) to plot at once and the default is set to 500 ms but can be increased to 1000 ms (1 second) depending on the available computing power. Data can be navigated using the scroll bar and by typing in the “Current Time” field the exact milliseconds at which a user wants to jump to. The other two text fields (Start Time and Stop Time) pertain to the start and stop times of an HFO event explained in detail later.

### 5. Visualizing Time-Frequency Representations

Time-frequency representation (TFR) of an event is important for determining if the event is an HFO. To visualize the TFR of a signal, a user must first click (indicated by a vertical black line) on any specific time point in the Main Window followed by opening the T-F Plots window (a button at the bottom of the Main Window). The top plot shows the raw trace of the selected source (users can choose the source in the Source drop-down menu at the top, it will only include sources that have been plotted on the Main Window). The middle plot is the filtered signal that can be modified by changing the “Low Freq(Hz)” and “High Freq(Hz)” parameters in the Filtered Signal Parameter section at the top of the T-F Plots Window. The bottom plot is the TFR centered around the time point that was selected on the Main Window. This graph is created with the use of the Stockwell Transformation. The warmer colors (red) represent frequencies at that time point where the power is the highest (thus contributes more to the morphology of the signal). The window size of this plot is 250 ms (this value can be changed via the “Window Size” parameter) and can be changed but the Stockwell transformation consumes a large amount of memory. The Stockwell Transform over-represents the higher frequencies, which makes it perfect for this application as the lower frequencies (that generally have high amplitudes) tend to drown out the higher frequencies.

The T-F Plots Window has a second tab called the “PSD Tab” which displays the Power Spectral Density (PSD) of the time point selected by the user in the Main window. One can think of the Stockwell Transform almost as an instantaneous PSD. The plotted PSD is just a slice of all the frequencies along that one timepoint in the Stockwell Transform’s output. This can be used to characterize the event as a Ripple (80 Hz - 250 Hz) or a Fast Ripple (250 Hz - 500 Hz).

### 6. Event scoring

Scoring of events is the main feature of hfoGUI tool. It allows users to find an event of interest (EOI) in their data and mark it for future visualization or analysis. The events can be saved after marking and can be loaded again for visualization or editing. Typically, when using this for HFOs, users would take these events, load them into their preferred coding language, extract features from these events, and use them to automate the process of identifying HFOs. To score the data, users must perform the following process:

a. Open the Score Window by pressing the “Score” button at the bottom of the Main Window, and a Score Window ((Figure 6)) will pop up.
b. To allow multiple users to score the data manually, there is a “Scorer” field that has to be filled out.
c. Now, users must select the Event of Interest time boundaries. To do this, users must proceed with the following protocol: Go to the Main Window. Press and hold the left mouse button starting at the beginning of the event. Drag the mouse to the end of the event, and release the left mouse button. Users should see that the desired data has now been highlighted which can be dragged from the ends of this highlighted section to fix the start or stop position. Users can also shift this highlighted section by clicking on the highlighted section, holding, and dragging the mouse.
d. Now that the temporal boundaries of the event have been selected, users can navigate back to the Score Window.
e. Select a score from the Score drop-down menu. The following score options are provided: Spike, Theta, Gamma Low, Gamma High, Ripple, Fast Ripple, Sharp Wave Ripple, Artifact, and Other.
f. Finally, users can press the “Add Score” button to confirm the score and will see this score now placed in the list located in the Score Window. The Score ID will be MAN[X], where MAN stands for Manual (if users manually curated this event). This is to distinguish between the manual and automated methods. Users can update the score by selecting them from the list, providing the correct score value, and pressing the “Update Selected Score” button at the bottom of the Score Window. If users select a score from this window, it will navigate the Main Window to this event. Therefore, users will not have to manually move the scrollbar to this point every time they want to show off their event of interest. If users decide that an event is no longer interesting, they can delete the score using a similar method to updating it. Select the desired score from the list, and then press the “Delete Selected Scores” button.
g. If users have found scores within this session that they want to save, they will need to press the “Save Scores” button towards the top of the Score Window. A pop-up window will have users provide a filename that they wish to save the scores as.
h. To re-load a session that users have previously analyzed, they can load the scores by opening the Score Window and selecting the “Load Scores” button. A pop-up will occur that will have them navigate to the score filename that users saved. After confirming the score filename, users will see the score list populate appropriately.

**Figure 6.**
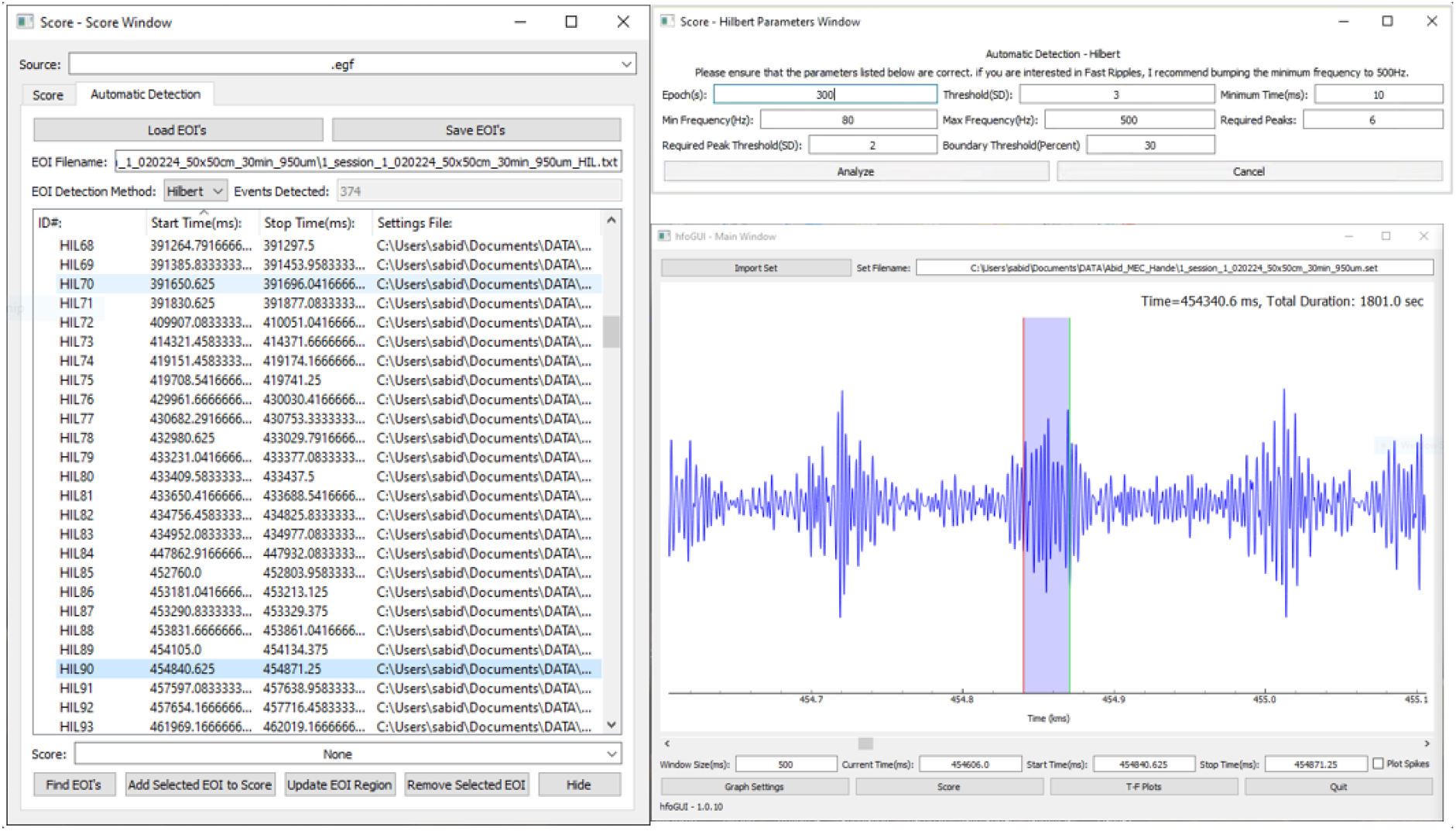
Score Window: *Left-* Clicking on “Score” button in the Main Window opens Scoring Window which has two tabs; Score and Automatic Detection. The Score tab (not shown) allows manual scoring of events by selecting (left mouse click, drag and release) an event in the Main window and assigning them as Theta, Gamma or any other frequencies from drop down menu. The Automatic Detection of events is done using Hilbert detection method and allows users to find an event of interest (EOI). The list shows multiple detected events after running the automatic detection. Columns show event ID, Start and Stop times of the events detected. HIL indicates detection using Hilbert method. Manually scored events would indicate MAN in the ID column. *Right top-* Users can enter different Hilbert parameters to achieve optimal detection based on the quality of the signal, experimental conditions etc. *Right bottom-* The main window shows the detected events has HIL90 event highlighted. The highlight shows Start (red line) and Stop (green) times which can be dragged from the ends manually adjust it as needed.

### Automatic HFO Detection

Manually scoring files can be time-consuming but there are a few automatic detection algorithms specifically for HFO detection, one of which we use is called Hilbert Detection [15]. To perform the automatic detection, users must first open the Score window, and navigate to the “Automatic Detection” tab. At the bottom of that window, users will see a “Find EOI’s” button. Once users click that, a pop-up window will allow them to provide the necessary parameters for automatic detection. Once users have finished changing the parameters, they may press the “Analyze” button to proceed with the automatic detection.

### Hilbert Detection

The detection algorithm implemented here is based on Burnos’s and RippleLab papers [16] [17]. We have listed the method below:

i. The data must first be band-pass filtered with the cutoffs at 80 Hz and 500 Hz, this will ensure that users are only including the HFO frequency bands. They use an Infinite Impulse Response (IIR) in the Burnos paper [16], but we use a 3rd order Finite Impulse Response (FIR), specifically a Butterworth filter. Their preference for an IIR filter over an FIR filter is due to the computational run time. We prefer FIR filters as to avoid phase delays introduced by IIR filters. If users prefer alternative cutoff frequencies, feel free to set the “Min Frequency (HZ)” and “Max Frequency (HZ)” parameters after pressing the “Find EOI’s” button.
ii. Analytic Signal / Envelope of the signal is calculated with the use of the Hilbert transformation.
iii. The envelope is binned into 5-minute (300-second) epochs. Users can change this 5-minute value by modifying the “Epoch(s)” parameter.
iv. The standard deviation of the envelope within the 5-minute bins is then calculated.
v. To find preliminary EOI’s a simple thresholding is applied. We suggest the mean of the envelope plus 3 standard deviations (within the bin). Users can modify this threshold parameter by changing the “Threshold (SD)” parameter.
vi. For each signal above threshold, the start/stop time must be determined. The Burnos paper determines this by when the signal reaches 50% of the max amplitude of the envelope of this event ((Figure 7)). The default in hfoGUI is 30% of the max but the “Boundary Threshold (Percent)” parameter can be modified as desired.
vii. These events are then discarded if they do not have a minimum duration of 6 ms. EOI’s with an inter-event-interval of less than 10 ms are merged together.
viii. It is here where the Burnos group diverts from the RippleLab group. The Burnos group then goes on to visualize the time-frequency representation of each of the events using a Stockwell Transform. This probably does a great job at discarding EOI’s that are unlikely to be an HFO, however, the Stockwell Transform tends to take a lot of time so hfoGUI uses more of a RippleLab approach from here. They still have the ability to visualize the Stockwell Transform of the signal, but it is not automated.
ix. The RippleLab group adds a secondary threshold and peak requirement. They require that at least 6 peaks exceed a threshold of mean + 3 SDs (they have their initial threshold set to 5 SDs). They can set the required peaks and the corresponding threshold by modifying the “Required Peaks” and “Required Peak Threshold (SD)” parameters respectively. They have 6 peaks above 2 SDs set as the default currently, but the user can modify these values. In our experience using a threshold (standard deviation or SD) of 4, required peaks of 6 and required peak threshold (SD) of 3 yielded the best results in terms of consistent ripples and fast ripples. It is advisable for the users to manually validate the events after running the automatic detection step since here are several factors that could affect the performance.

**Figure 7.**
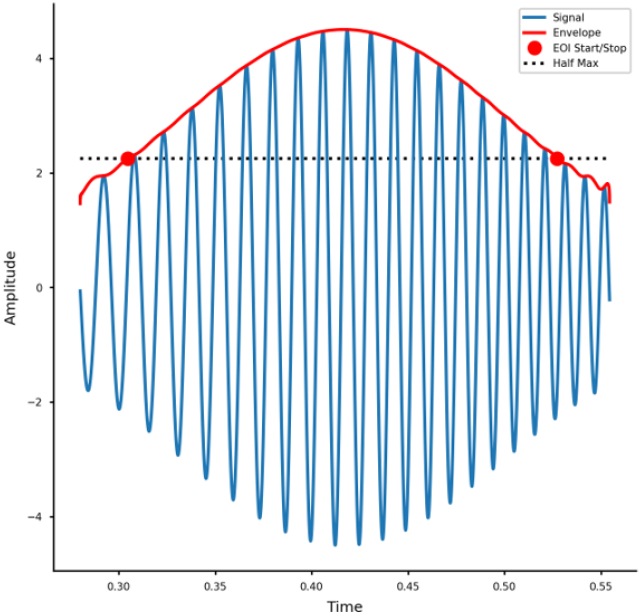
Hilbert transformation of a signal (blue), with envelop (red) and half max at 50% (dotted line). The two red dots indicate the start and stop points of half maximum amplitude.

### SSM— Visualization and Spatial Analysis of LFPs

The majority of neurological research employing electroencephalography (EEG) focus on select frequency bands; Theta, Beta, Delta, and Gamma because they are physiologically significant oscillations in mediating the encoding/decoding mechanisms of memory consolidation and retrieval [18–20]. However, they generally focus only on temporal changes in spectral density mapped back to brain regions during behavior experiments. Since “spatial mapping” of signals primarily relates to mapping spectral information to regions of the brain a tool to correlate frequencies to subject’s physical position is sorely missing. In spatial navigation research, scientists are interested in how brain oscillations are influenced by subject’s behavior while navigating an environment. Spatial spectral mapper (SSM) fills this gap by providing a way to visualize and analyze frequencies combined with subject’s positional data. The tool provides a heatmap of overall frequencies with respect to specific time bin the animal occupied specific positions in the arena. For example, one can visualize the power of theta signal for the entire session in 5 second chunks to investigate instantaneous actions and understand the ongoing brain oscillations during specific behavioral states. For instance, it can be used to investigate how specific frequencies change with certain behaviors such as grooming, thigmotaxis, exploration/attention (objects, odors) etc. [21–25]. Studies have implicated specific frequencies to dominate during navigational or attentional tasks and this tool will help unravel the role of neural oscillations during specific behavioral modes [26, 27].

### Installation and Usage

SSM was built using Python and was specifically tested using Python 3.8. There is no Operating System requirement as SSM utilizes the PyQt5 framework (for GUI support) which is cross-platform. However, this tool was tested using Windows 10. SSM is available from github.com/HussainiLab/SpatialSpectralMapper and installation details are described there. It currently takes data produced in the Axona recording format but we provide a tool below to allow conversion of other data to Axona format. SSM has a main window (Figure 8) where user loads a file ‘.eeg’ or ‘.egf’ file for the EEG recording, and a ‘.pos’ file for subject tracking. SSM offers support for 7 well-documented frequency bands; Delta (1-3 Hz), Theta (4-12 Hz), Beta (13-20 Hz), Low Gamma (35-55 Hz), High Gamma (65-120 Hz), Ripple (80-250 Hz) and Fast ripple (250-600 Hz). SSM’s chunk size widget and slider offer a choice to the users to chunk EEG data into ‘n’ second intervals in the time domain prior to PSD mapping. SSM also supports printing time series PSD data directly into an Excel file using its save data feature.

**Figure 8.**
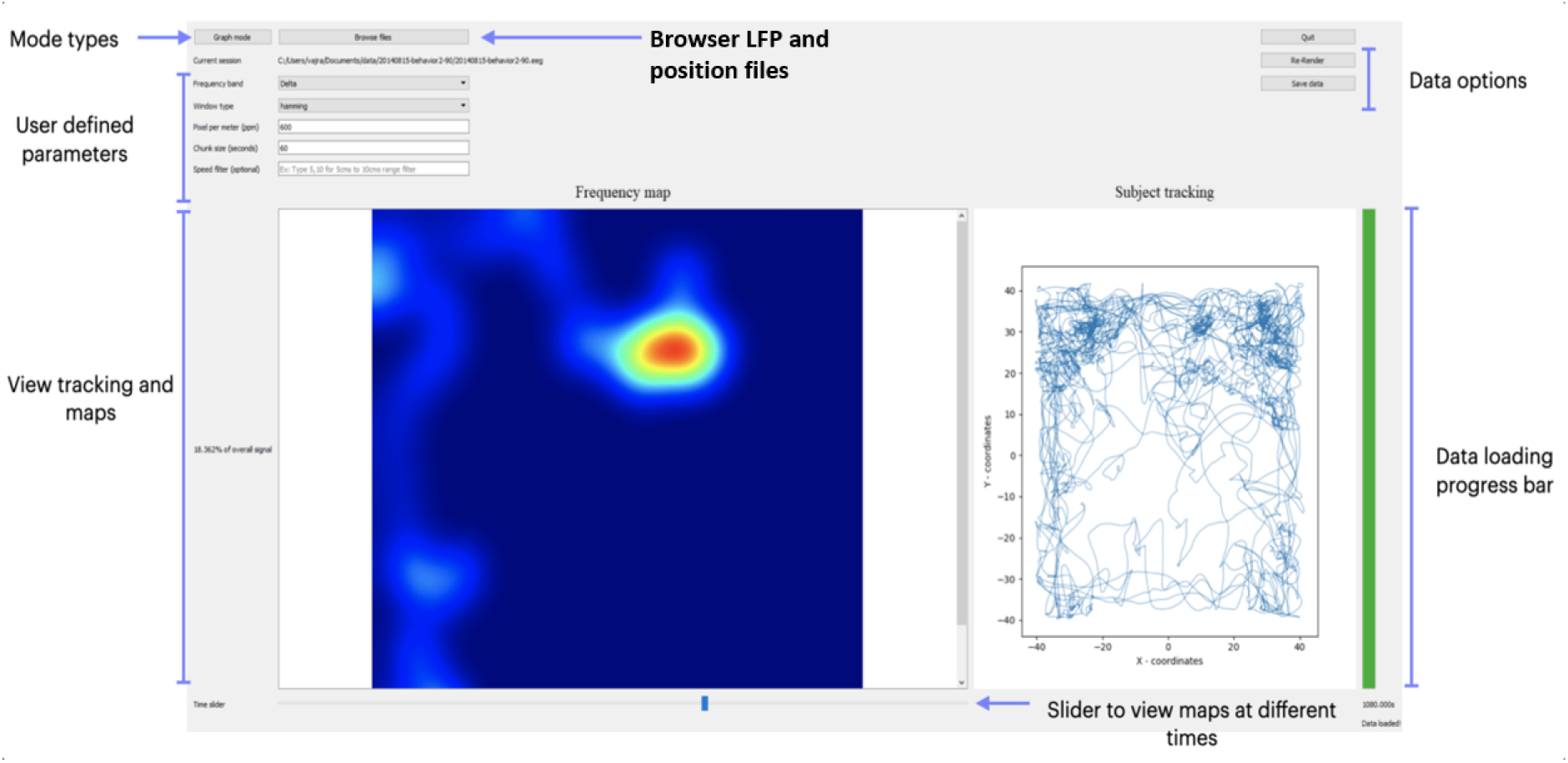
Spatial Spectral Mapper (SSM): The core feature of the SSM tool is the *Frequency Map* which displays the power spectral densities (PSDs) of an LFP signal as a spatial heatmap indicating low (blue) to high (red) PSD. The slider at the bottom allows visualization of the maps at different time range depending on the *Chunk Size* set by the user. Other parameters such as *Window type* and *Speed filter* affect the frequency maps too. The *Subject tracking* window on the right reflects cumulative arena coverage of the animal at each time interval. The “Graph mode” button allows visualization of PSD graphs from the Welch computation instead of frequency maps (see Figure 9). The “Save data” button saves frequency PSDs to an Excel file.

### Steps for using SSM

1. Users must define specific parameters before choosing and analyzing LFP data (Figure 8). First, the user must choose a window type for the Welch computation. Options include Hamming, Hann, Blackman, and boxcar window. Default is Hamming window.
2. Users must then set the Pixel Per Meter (PPM) value based on their arena dimensions for position tracking. This value helps scale how many pixels of movement correspond to a physical distance in the ‘.pos’ file. This value can be calculated easily if the dimensions of the physical maze and camera resolution are known.
3. The chunk size is set in an integer number of seconds. Chunk size determines how the LFP signal is split up. For example, a chunk size of 5 seconds implies the LFP signal will be split into 5 second intervals, and a PSD is computed for each 5 seconds of signal.
4. Users have the option to set a speed filter range if needed. This allows the user to filter out any position tracking data that falls within a specified range. For instance, an input of “5, 30” will ensure that only tracking data between 5cms and 30cms is included for mapping.
5. Users than have to select ‘Browse Files’ and navigate to a directory containing ‘.eeg’ and ‘.pos’ file types. These file types are typically produced by the Axona recording system but other file formats can be converted to Axona format and used here. Once both ‘.eeg’ and ‘.pos’ files from the same session are picked and OKed, map computations will immediately begin as indicated by the progress bar.
6. Once the computation is done, a complete set of maps for all frequencies are computed and the user can simply choose a specific frequency band to check their percent power on the left of the GUI.
7. If the PPM, window type, chunk size, or speed filter options are changed, users have to re-compute the set of frequency maps with these new settings. The ‘Re-render’ button will turn green to indicate this and upon clicking it will generate a new set of frequency maps with the altered settings.
8. The user can scroll through the frequency maps using the time slider at the bottom of the window, to visualize how the frequency maps evolve over the course of the experiment. A tracking window on the right reflects the subject’s cumulative arena coverage at each time interval.
9. Users can click the ‘Graph mode’ button to view the set of PSD graphs from the Welch computation in place of the frequency maps for each time interval (Figure 9).
10. Finally, the ‘Save data’ button will export all frequency PSDs for each time chunk to an excel file for further analysis.

**Figure 9.**
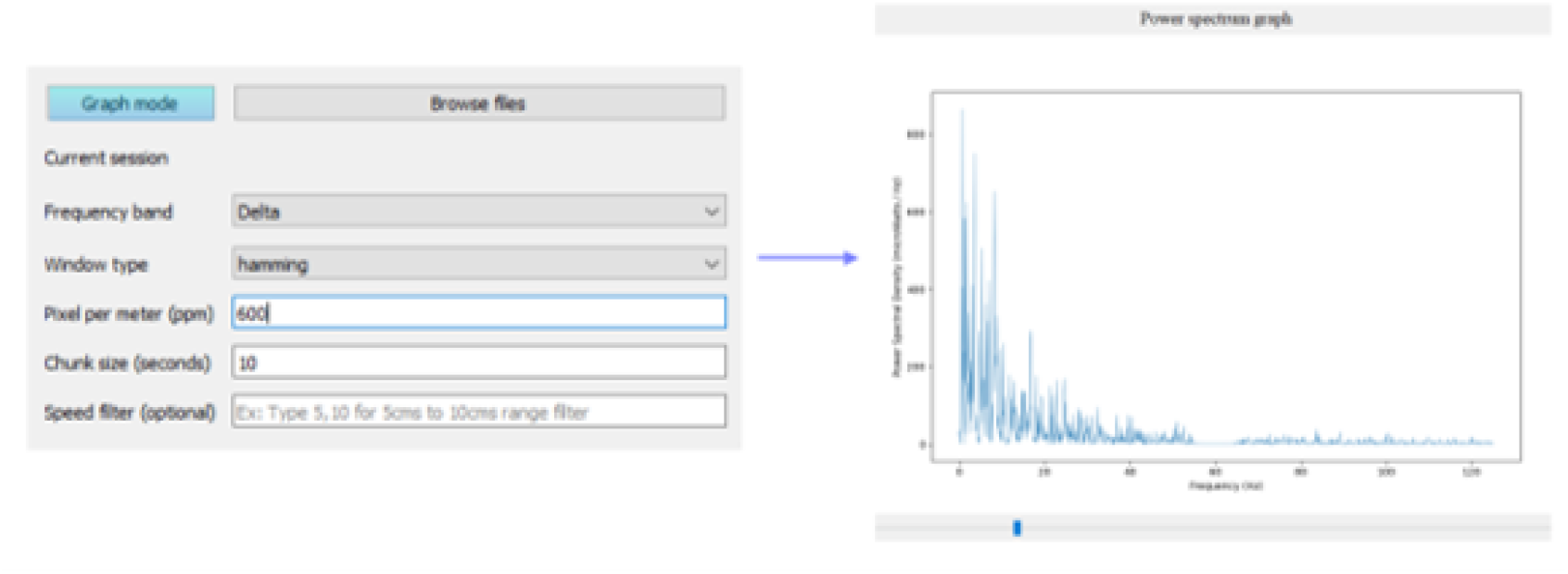
Graph mode (PSD): The graph mode displays the power spectral density of the signal at specific time (chunk) range. This is useful to quickly check which frequencies are dominating at a specific location in the arena.

Note that a very small chunk size such as 1 or 2 seconds would take a long time to compute maps.

### Data types and Converter

Both hfoGUI and SSM tools only take in Axona data format files (.eeg and .egf) but we provide a converter that allows conversion of Intan (.rhd) files to Axona file format. Any position data in form of .csv file will also be converted to Axona’s position file format (.pos) which is required for using SSM. The converter is available here: https://github.com/HussainiLab/intan_to_axona/

## Discussion

The development of hfoGUI and SSM addresses key limitations and advances capabilities within open-source LFP analysis software for the neuroscience community. We demonstrate their specific strengths in facilitating the investigation of both temporal and spatial dynamics inherent to local field potential data in rodent models.

Research into high-frequency oscillations (HFOs) has accelerated in recent years due to their emerging role in neural information processing and potential as biomarkers for neurological disorders. Our hfoGUI tool directly contributes to this vital research area by providing an accessible and specialized platform for HFO analysis. Unlike many existing tools that depend on proprietary platforms, hfoGUI leverages the Python ecosystem and provides a user-friendly graphical interface, expanding its reach within the research community. hfoGUI’s optimized implementation of the Hilbert detection method for HFOs represents a significant innovation. Drawing inspiration from the work of Burnos et al. (2014) and RippleLab [17], we strike a balance between computational rigor and efficiency. While the Stockwell transformation offers valuable insights, our focus on signal peak thresholds and peak number criteria accelerates HFO detection without sacrificing accuracy. This refinement makes hfoGUI suitable for rapid analysis of large datasets, facilitating the exploration of HFOs in diverse behavioral and neurological contexts.

The SSM tool opens a new dimension in the understanding of spatiotemporal relationships within LFP/EEG data. To our knowledge, SSM is the first tool in its ability to provide an open-source, interactive platform to process and visualize spectral signal encoded as heat map information based on subject’s position. The ability to analyze spectral power distribution allows for targeted investigation of how neural oscillations underpin specific behaviors such as spatial organization of theta oscillations during navigation or gamma oscillations during attentional tasks [26, 27]. Such analyses hold the potential to elucidate the neural mechanisms governing cognition, memory, and complex behaviors.

The availability of hfoGUI and SSM offers advantages and could lead to further innovation in neuroscience research. Reliance on proprietary software can constrain research, especially for early-career scientists and laboratories with small budgets. By providing free and open-source tools, we foster collaboration, reproducibility, and broader participation in LFP research. The focus on temporal HFO analysis and spatial LFP mapping addresses specific research needs that are often under-supported by general-purpose analysis software. There are however caveats to using open-source compared to paid software that come with customer support. Open-source solutions offer flexibility and customization but users may need a degree of technical expertise to fully harness their potential. Additionally, researchers must assume full responsibility for validating the results obtained using hfoGUI and SSM. Some other caveats include lack of batch processing options to run multiple files or folders and limited options to import different data types at this time.

In the future we envision incorporating machine learning algorithms to automate HFO detection and classification, enabling even larger-scale studies and the identification of subtle patterns. Adapting hfoGUI and SSM for real-time LFP processing would unlock opportunities for innovative closed-loop experiments, neurofeedback paradigms, and the development of advanced brain-computer interfaces.

With hfoGUI and SSM, we demonstrate the value of open-source solutions for advancing neuroscience research. By addressing specific analysis needs, promoting accessibility, and creating a foundation for collaboration, these tools contribute to a more inclusive, efficient, and transformative research community.

## Author contributions

G.M.B. and S.V. were the major contributors for hfoGUI and SSM tools respectively. G.M.B., S.V., and S.A.H. conceived the idea, discussed the methods and results of the tools. G.M.B. and S.V. wrote the manuals for hfoGUI and SSM tools respectively, and S.A.H. wrote the manuscript. O.S. wrote the code for the file converter with help from A.A. S.A.H. supervised the project and obtained the funding.

## Acknowledgements

We would like to thank Dr. Alexandra Petrache, Nikhil Ramavenkat, Sandhya Senthilkumar, Matthew Hou, Jessica George and Satya Baliga for testing various parameters of hfoGUI. We also thank Dr. Gustavo Rodriguez and Dr. Radha Raghuraman for providing their data for visualization purposes on this manuscript. These tools were supported by NIH/NIA grants R01AG050425 and R01AG064066.

## Notes

### Competing Interest Statement

The authors have declared no competing interest.

